# Fighting over food unites the birds of North America in a continental dominance hierarchy

**DOI:** 10.1101/104133

**Authors:** Eliot T. Miller, David N. Bonter, Charles Eldermire, Benjamin G. Freeman, Emma I. Greig, Luke J. Harmon, Curtis Lisle, Wesley M. Hochachka

## Abstract

The study of aggressive interactions between species has, to date, usually been restricted to interactions among small numbers of ecologically close competitors. Nothing is known about interspecific dominance hierarchies that include numerous, ecologically varied species. Such hierarchies are of interest because they could be used to address a variety of research questions, e.g. do similarly ranked species tend to avoid each other in time or space, and what will happen when such species come into contact as climates change? Here, we propose a method for creating a continental-scale hierarchy, and we make initial analyses based on this hierarchy. We quantified the extent to which a dominance hierarchy of feeder birds was linear, as intransitivities can promote local species’ coexistence. Using the existing network of citizen scientists participating in Project FeederWatch, we collected the data with which to create a continent-spanning interspecific dominance hierarchy that included species that do not currently have overlapping geographic distributions. Overall, the hierarchy was nearly linear, and largely predicted by body mass, although there were clade-specific deviations from the average mass–dominance relationship. Most of the small number of intransitive relationships in the hierarchy were based on small samples of observations. Few observations were made of interactions between close relatives and ecological competitors like *Melanerpes* woodpeckers and chickadees, as such species often have only marginally overlapping geographic distributions. Yet, these species’ ranks—emergent properties of the interaction network—were usually in agreement with published literature on dominance relationships between them. *Interspecific dominance hierarchy, aggression, displacement, citizen science*

**LAY SUMMARY:** When it comes to fighting over food, bigger is better but woodpeckers are best. The outcome of aggressive encounters between birds frequently determines which individual gains access to contested resources like food, but until now, little was known about such encounters between individuals of different species. We partnered with citizen scientists to record interspecific behavioral interactions at bird feeders around North America, and assembled these interactions into a continental dominance hierarchy.

## INTRODUCTION

Many animals congregate at food resources that can be patchily abundant. Examples include carcasses on the Serengeti Plain and bird feeders in suburban St. Louis, USA. Frequently, individual animals maintain access to the resource by physically dominating others. When the direction of aggression between individuals of different species is consistent, an interspecific dominance hierarchy is created (Drews 1993), with important consequences for access to resources (LeBrun 2005). Dominance hierarchies are more often studied at the intraspecific level, where previous work has focused on social interactions between individual animals (Farine and Whitehead 2015). Of the studies on interspecific social dominance, those hierarchies studied to date typically have included only a small number of closely related and/or ecologically similar species (Morse 1974; Wallace and Temple 1987). Importantly, studies at the interspecific level have tended to focus on competitive rather than social dominance (Syme 1974). Competitive dominance is when, under some set of particular environmental conditions, individuals of one species have higher fitness than individuals of another species (Zamudio and Sinervo 2000). Social interactions, the purview of this paper, are not necessarily relevant to competitive dominance, but nevertheless can have major impacts on communities. For example, in Australia the socially-dominant Yellow-throated Miner (*Manorina flavigula*) aggressively ousts most other co-occurring passerine birds to such a degree that the miner’s presence demonstrably reduces other species’ abundances (Kutt et al. 2016). Similar interspecific aggression by Bell Miners (*M. melanophrys*), another aggressive Australian bird, initiates a trophic cascade wherein exclusion of insectivorous passerines by Bell Miners leads to local infestations by tree sap-feeding psyllids (Hemiptera), and these high-density psyllids ultimately cause the local decline of multiple *Eucalyptus* species (Loyn et al. 1983). Aggressive interactions may even influence continental-scale patterns of occupancy; for example, dominant bird species arrive earlier and migrate shorter distances (Freshwater et al. 2014). In spite of their potential importance, even basic information on the structure and correlates of interspecific social dominance hierarchies is typically lacking.

Dominance hierarchies can take many structural forms (Fig. 1), from perfectly linear pecking orders (Perrin 1955) to despotic systems, where one species (or individual in the traditional intraspecific context) is dominant and all others are subordinate, and to corporative systems, where a ranking order exists, but some species hold equivalent ranks (Fushing et al. 2011). With respect to interspecific competitive dominance, a current research focus is the degree to which such hierarchies deviate from perfect linearity, as deviations from linearity can facilitate species coexistence (Gilpin 1975; Laird et al. 2006; Allesina and Levine 2011). Departures from linearity manifest themselves in the form of mathematical intransitivities. An example of an intransitivity is when species A is dominant to species B, B is dominant to C, but C is dominant to A. Known as a rock-paper-scissors relationship in behavioral ecology (Zamudio and Sinervo 2000), such a relationship indicates that despite pairwise competitive advantages, no species is able to gain preferential access to resources over all species, and therefore no single species can exclude all others. Network theory provides a framework to quantify the prevalence of intransitive relationships within those hierarchies (Shizuka and McDonald 2012), as well as the structure of dominance hierarchies (Farine and Whitehead 2015) and species’ positions therein.

**Figure 1.**
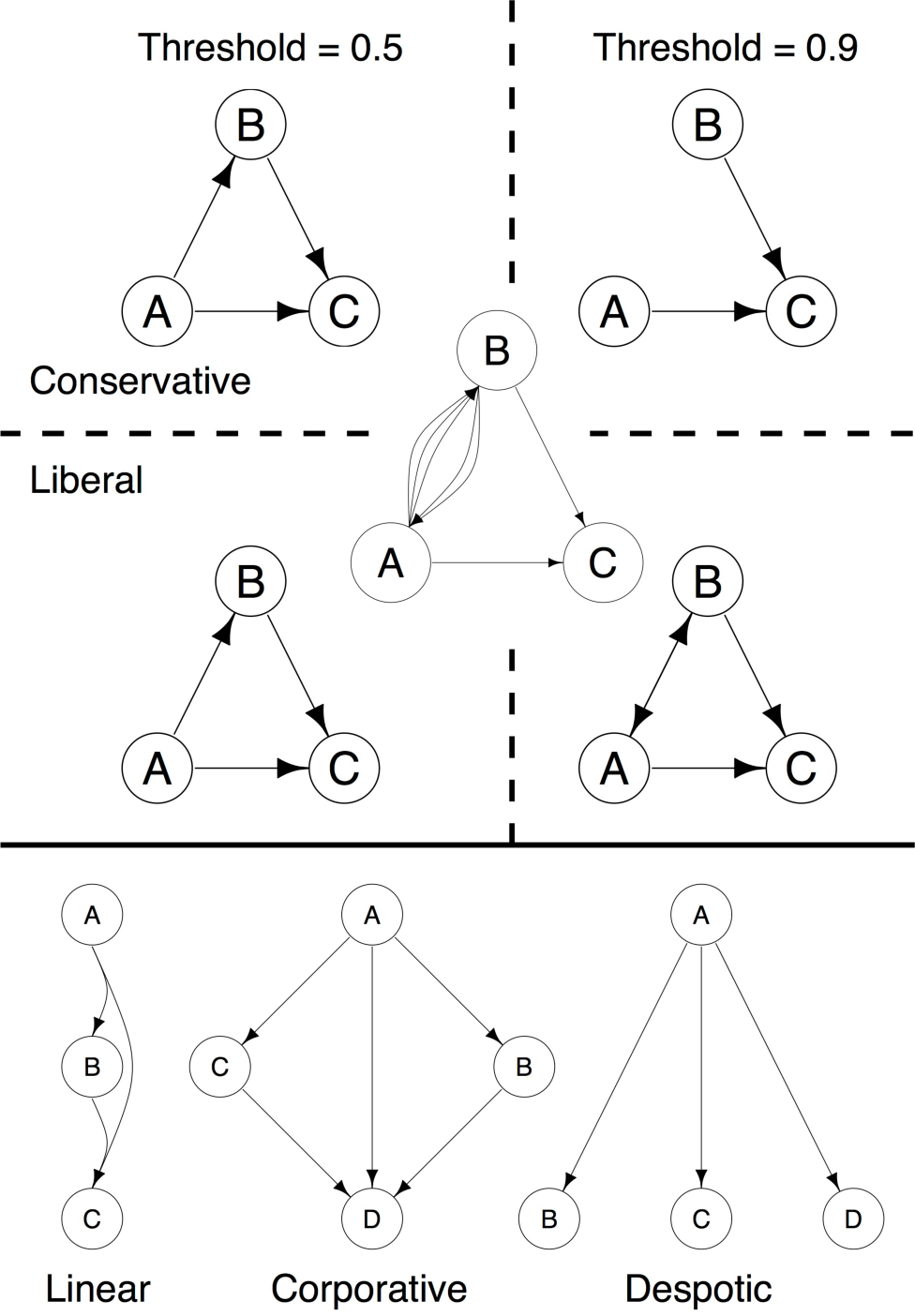
The upper panel uses a series of schematics to explain the network simplification step of the directed acyclic graph (DAG) method of identifying intransitivities in the continental interspecific dominance hierarchy. Given an input of seven unique observations like those represented by the middle subnetwork, we converted the raw observations into networks where interactions between species were either absent, unidirectional, or bidirectional. For each pairwise interaction (e.g., A vs. B), if one of the species won more than the threshold proportion of wins (0.5 in the left-hand subnetworks, 0.9 in the right-hand side), then we collapsed all A vs. B interactions down to a unidirectional interaction running from the dominant to the subordinate species. If neither species won more than the threshold, then we either removed all A vs. B interactions (conservative DAG, top row of subnetworks), or set A vs. B to a bidirectional interaction (liberal DAG, bottom row of subnetworks). We then detected intransitivities in the subnetworks by determining whether the graph was directed acyclic or not, with the caveat that not all cyclic graphs are necessarily intransitive with the liberal DAG method. For instance, the bottom right subnetwork contains a cycle, but A = B > C is not an intransitive relationship. The lower panel shows a series of example dominance hierarchies of different forms.

Studies of relatively small taxonomic scope have suggested that species’ traits, particularly body mass, are important predictors of a species social dominance in pairwise interactions of species. Martin and Ghalambor (2014) showed that in pairwise interactions, socially-dominant species were typically larger. There also appears to be variation at larger taxonomic scales (i.e., between genera or families), although the traits in question, be they behavioral or morphological, have rarely been identified. In one such rare study, among the four bird families studied by Martin and Ghalambor (2014), some groups, e.g., those that foraged on trunks, were less dominant than expected based on their body mass. Still unknown is whether clade-level determinants of interspecific social dominance overwhelm the signal of body mass in structuring dominance hierarchies at broader taxonomic scales.

The structure of interspecific dominance hierarchies remains understudied in part because of the inherent difficulty of compiling sufficient behavioral observations to describe a hierarchy, though some work has been done with hummingbirds (Feinsinger 1976; Wolf et al. 1976), and research into the ramifications of intraspecific dominance hierarchies is also of relevance (Hobson and DeDeo 2015). Gathering behavioral data at a large scale can be achieved by partnering with citizen science initiatives that engage the public in protocol-driven research. Because supplemental feeding of birds concentrates birds at high density and in a context that facilitates the logging of standardized behavioral observations, bird feeders provide an excellent venue at which to observe interspecific interactions. We collaborated with citizen scientists to catalog interspecific aggressive interactions (displacements) between birds at feeders, and used these behavioral observations, collected across the breadth of the North American continent, to rank numerous, distantly related species in a common framework. We used this dominance hierarchy to address two hypotheses: (1) feeder-attending birds follow a predictable and roughly transitive “pecking-order” dominance hierarchy across a broad spatial extent; and (2) body size mediates species’ positions within this dominance hierarchy, with heavier species more dominant than lighter species. We quantify deviations from perfect linearity, and identify the species involved in these relationships.

## METHODS

### Data collection

Aggressive interactions between birds can take several forms. In this study, we focused on the most overt of these: displacements in which one bird displaces or chases another from a perch at or near a bird feeder. We did not include subtle interactions such as one bird waiting until another finished feeding, or accidental displacements such as one bird arriving suddenly and temporarily frightening off birds that had been present.

To collect these behavioral data, we partnered with Project FeederWatch (PFW). PFW engages more than 20,000 people annually to monitor the abundance and distribution of feeder birds in the U.S. and Canada (Bonter and Cooper 2012) during the non-breeding season (early November until early April). Participants follow a standard observation protocol and submit counts of the maximum number of individuals of each species that are simultaneously using their feeder over a two-day period. Beginning in February 2016, we augmented the data input protocol to allow participants to report displacements observed during their counts. Data from PFW, including the new behavioral interactions, are freely available online (http://feederwatch.org/explore/raw-dataset-requests/), and the interactions database continues to grow by approximately 110 observations per day (as of January 2017). Currently, ~11% of FeederWatchers have augmented their checklists with behavioral data. The database used in the analysis of this paper contained 5,986 observations submitted prior to 27 December 2016, of which 3,616 were interspecific displacements.

Many participants in PFW already exhibit a high level of natural history knowledge. Nevertheless, to standardize observations among participants, we used the online PFW blog to communicate to participants the behaviors of interest and provide accompanying example videos (http://feederwatch.org/uncategorized/different-bird-behaviors-explained/). To further quality-check our database, we both manually monitored incoming observations, and wrote R scripts to automatically flag unusual interactions. Examples of such observations were those involving rarely seen birds, or between species where previous evidence would suggest the reported direction of aggression would be rare. Approximately 15% of observations were flagged during this initial stage, but many of these observations were manually approved by FeederWatch staff (i.e., with no further contact with the participant) after reading through comments provided by participants. Participants responsible for observations still flagged after this point were contacted directly (approximately 2% of all observations). If the participant responded with details confirming the observation, we removed the flag and the observation was included in the data used in this paper. Prior to November 2016, participants were unable to delete their submitted interactions. Flagged, unconfirmed observations from before this date were removed from the database by the authors (approximately 2% of total data at that time). Participants are now able to delete and correctly re-enter observations, so flagged, unconfirmed observations are simply excluded from analysis, rather than removed from the database (less than 1% of total observations). We also excluded from analyses observations involving potential predators and vultures, as interactions with potential predators and vultures can be difficult to categorize, and between Broad-billed and Costa’s Hummingbirds, as these hummingbird species were only observed to interact with one another.

### Network creation and calculation of rank

We used the 3,616 interspecific observations of displacement to create a directed, weighted network. Each unique interspecific interaction was represented by a directed edge (an arrow) between nodes (species). We developed and used in this paper a modified form of Bradley-Terry model to rank species in a hierarchy. The original Bradley-Terry model fits a function wherein all species are ranked against a single focal species (Turner and Firth 2012). In preliminary analyses we found that Bradley-Terry models (Bradley and Terry 1952) returned a ranking in general agreement with published pairwise interactions (Rodewald 2015). Our modification was to consider each species in turn as the focal species, and then take species’ median coefficients across all fitted models as their overall dominance scores. We sorted these and assigned ranks accordingly. To confirm that our results were not sensitive to inclusion of species with limited information, we repeated all analyses after excluding species for which fewer than ten interactions were observed, and we also repeated all analyses restricting observations to the eastern United States, our area of highest data density. Results were qualitatively identical, so we present results from the complete dataset here.

We had evaluated and eliminated from consideration other approaches to creating a dominance hierarchy. We did not use eigenvector centrality and Elo score rankings as both artificially inflated the ranks of species that had interacted a limited number of times, but that when observed had interacted with high-ranked species. For example, based solely on the fact that it had lost to the fairly high-ranked House Finch (*Haemorhous mexicanus*) and White-crowned Sparrow (*Zonotrichia leucophrys*), Mountain Chickadee (*Poecile gambeli*) was consistently ranked higher than Black-capped Chickadee (*P. atricapillus*), a species to which Mountain Chickadee is known to be subordinate to (Minock 1972).

### Phylogenetic signal of dominance and relation to body size

As a test of whether social dominance is a trait that exhibits phylogenetic signal, we calculated Pagel’s lambda (Pagel 1999) for species’ Bradley-Terry model coefficients. Pagel’s lambda varies from zero to one, where one indicates that variance in dominance is well predicted by a Brownian motion model of evolution, i.e., phylogenetic covariance predicts dominance. That said, signal in dominance rank detected here might be a byproduct of body mass, which is well known to exhibit strong phylogenetic signal (Blomberg et al. 2003). Moreover, clade-level deviations from the overall dominance-mass relationship are potentially interesting. To address these issues, we tested the degree to which body mass can explain the variance in species’ Bradley-Terry model coefficients with a phylogenetic generalized least squares (PGLS) regression (Grafen 1989; Martins and Hansen 1997), implemented in the R package *caper*. We used the maximum-likelihood optimized lambda in the model (a measure of the phylogenetic covariance of the residuals in dominance rank), log-transformed species’ average body masses as published in Dunning (2007), and a pruned version of the maximum clade credibility global bird tree (Jetz et al. 2012). We examined species’ residuals from the fitted PGLS to identify clades with notably positive or negative dominance scores given their body mass.

### Calculation of network transitivity

If we have a dominance hierarchy where pairwise relationships are represented by a single direction of aggression, and it is still possible to trace a route from one species back to itself in that network, then it contains at least one intransitivity of the sort thought to promote species coexistence (Allesina and Levine 2011). Put differently, such a situation arises when all pairwise species interactions are consistent, and there are no contested relationships where sometimes species A wins and other times B wins, but it is still possible to follow directed aggression through the network and return to the same player. In network theory jargon, if a network has no parallel and/or bidirectional edges, and if it is not a directed acyclic graph, then somewhere in that dominance hierarchy an intransitive relationship exists.

At a three-species level, we can visualize this as a biological version of rock-paper-scissors, but intransitivity can manifest itself across larger numbers of species. Quantifying the degree of intransitivity in a network with more than a few nodes (species) is computationally challenging. Most algorithms work by assessing all possible subnetworks of a given size within the actual dominance hierarchy. For instance, with 112 species (the number included in our study), there are 6,216 possible 2-species networks to consider, 227,920 possible 3-species networks to consider, and 3.9^32^ possible 56-species networks to consider. Because it has an empirical solution, we employed the triangle transitivity method (Shizuka and McDonald 2012), which works by calculating the proportion of transitive triangles (three-species subnetworks) relative to all triangles. This measure, *P_t_*, can be thought of as a measure of how many rock-paper-scissors relationships there are within the larger network. The triangle transitivity method has limitations, however: *P_t_* is assessed at the triad level, with no tests for intransitivity within groups of four or more species, and it simplifies two-species interactions according to a majority rule. For example, if two participants observed a Downy Woodpecker (*Picoides pubescens*) displace a Red-breasted Nuthatch (*Sitta canadensis*), but one observed the opposite direction of aggression, the triad approach would simplify this to Downy Woodpecker as dominant to Red-breasted Nuthatch. This approach does not consider the transitivity of triads where one relationship is unknown (e.g., “pass-along” relationships *sensu* Shizuka and McDonald 2012).

To probe some of the complexity lost to the triangle transitivity method, we additionally developed a directed acyclic graph (DAG) method. We used the DAG method to determine how our calculations of intransitivity varied over two-, three-, and four-species subnetworks, and over variable thresholds (proportions of wins) a species needed to have over a competitor before it was considered dominant. After initial exploration of how the degree of intransitivity varied across different values, for the two-and three-species subnetworks, we chose to examine the following thresholds of percentages of wins by the dominant competitor: 50, 55, 60, 70, 80, 90 and 100%. For the four-species subnetworks we examined the thresholds: 50, 60, 80, and 100%. We also explored the sensitivity of the intransitivity results to how ties were handled. When neither individual in a pairwise interaction won more than the threshold proportion of interactions, we either removed the interaction entirely or set it to a tie (i.e., created a bidirectional edge where A = B). The first of these approaches, which we refer to as the “conservative DAG” method, is a more conservative way of identifying biologically relevant intransitivities. Any subnetwork identified as not being a DAG here is indeed intransitive. The same is not necessarily true for the second approach (Fig. 1). Despite this, we chose to explore this “liberal DAG” method, since it maintains a larger proportion of the data. The DAG methods assess the transitivity of pass-along and other relationships where all possible links within the subnetwork are not necessarily known.

We identified which species tended to be involved in these intransitive relationships by deriving species-specific standardized effect sizes (SES) that reflected the proportion of intransitive relationships in which species were involved, compared with expectations. To do this, we focused on the three-species, 50% threshold, conservative DAG method’s results. We compared the observed number of intransitive relationships in which each species was involved to the expected number based on 999 runs where we randomized the winner and loser of every observation in the input data, and summarized the resulting network with the conservative DAG method. Large negative values of this measure reflect species that, after accounting for prevalence in the dataset, were involved in few intransitive relationships relative to expectations, while large positive values reflect species that were involved in more transitive relationships than expected by chance.

## RESULTS

### Network creation, calculation of rank, and relationships to body mass

We sorted and ranked species’ median Bradley-Terry model coefficients to produce the hierarchy illustrated in Fig. 2. Pagel’s lambda, a measure of the phylogenetic signal in dominance, was 0.81, and was significantly different from zero (*p* = 0.001). Body mass explained some of the variance in species’ coefficients from the fitted models (PGLS pseudo-*R*^2^ = 0.31, *p* < 0.001). The optimized PGLS lambda value of 10^−6^, however, illustrates very limited phylogenetic covariance in the residuals (distinct from the explanatory power it has with respect to species’ dominance scores *per se*). Taken together, these results show that, on average, larger bird species are more dominant, but that dominance is affected by other factors that render entire clades (e.g., families, Fig. 3) more or less dominant than the average; additionally, there is substantial variance in dominance that remains unexplained. As examples of these clade-specific effects, we found that woodpeckers tended to be more dominant than expected based on their body mass, while other lineages such as chickadees and titmice tended to be less dominant than expected based on body mass (Fig. 3).

**Figure 2.**
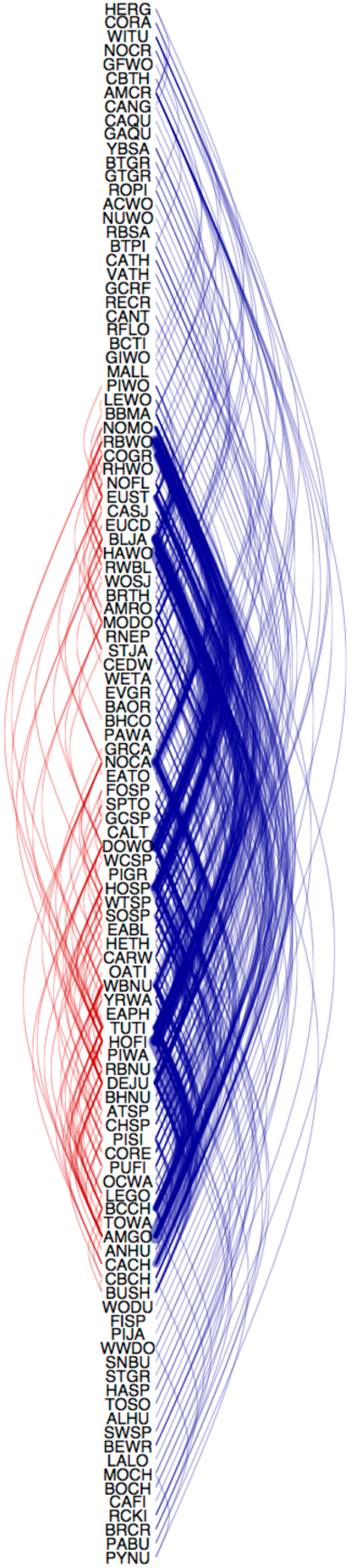
Attribute-ordered network showing the species ranked from most dominant (top) to most subordinate (bottom) according to their Bradley-Terry model coefficients. Blue lines represent observed displacements from an inferred dominant to an inferred subordinate species, while red lines represent observations where a species with an inferred lower dominance rank displaced one with an inferred higher rank. The preponderance of blue lines emphasizes that most observed interactions were in the direction that would be expected given a linear dominance hierarchy.

**Figure 3.**
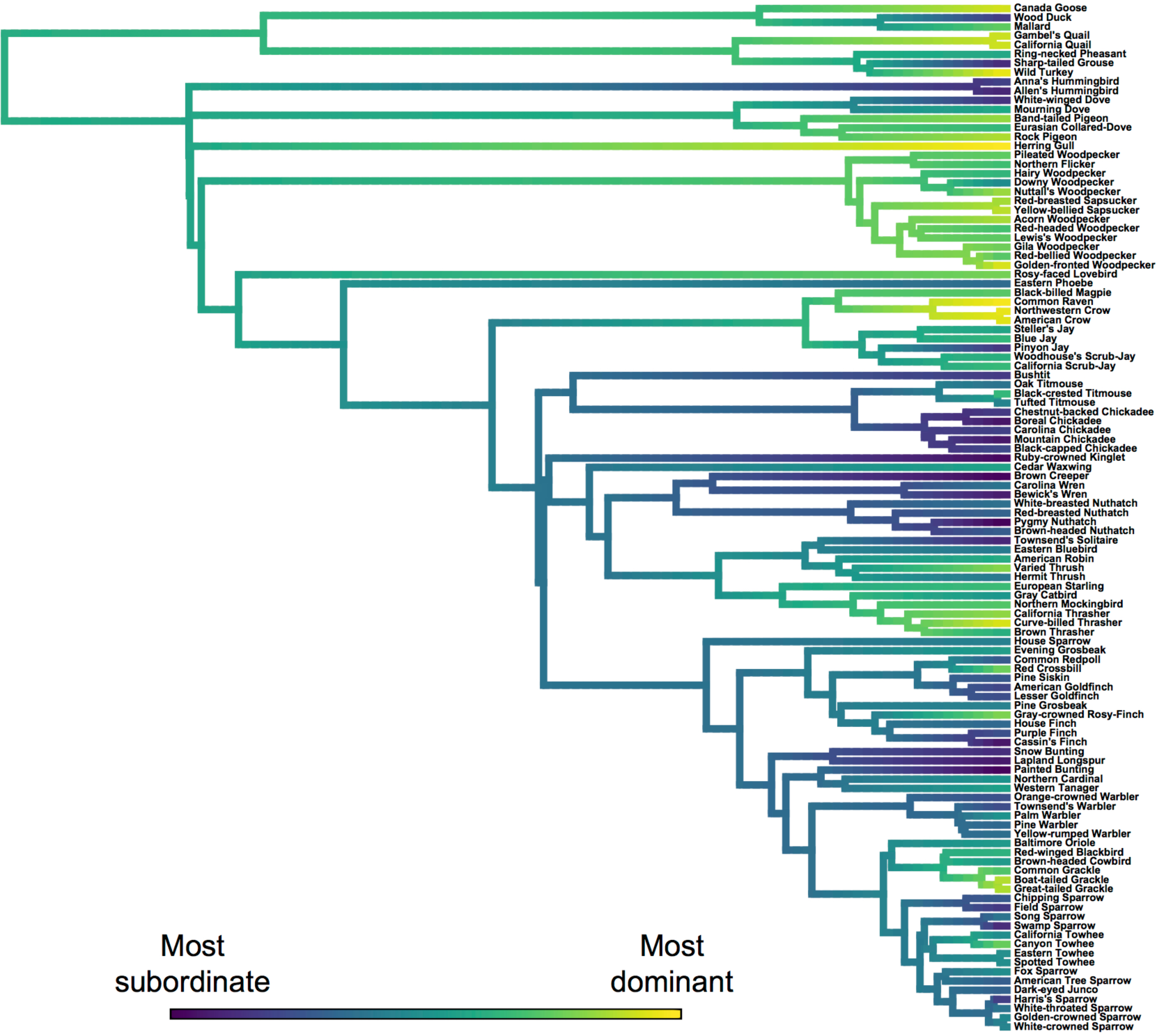
Dominance rank color-coded on the bird phylogeny. More dominant species are colored in yellow, while more subordinate species are colored in blue. The low rank of chickadees, titmice, nuthatches and wrens, can be seen here, as can the comparatively high rank of woodpeckers, jays, crows, and ravens.

### Calculation of network transitivity

According to the triangle transitivity method (Shizuka and McDonald 2012), the continental interspecific dominance hierarchy was more linear than expected based on chance (*P_t_* = 0.98, *t_tri_* = 0.92, *p* < 0.001). Put differently, 98% of all known possible three-species subnetworks were transitive. Some of the pairwise interactions that generated intransitivities can be seen as red lines in Fig. 2.

We developed the directed acylic graph (DAG) method to further examine subtleties in species’ interactions. Here, we considered two-, three- and four-species subnetworks, various thresholds (proportions of wins) a species needed to attain to be named the dominant member of a two-species interaction, and we considered two alternative methods of handling tied two-species interactions, where a tie was defined as neither member of a species-species interaction winning more than the threshold proportion of observed interactions. Across all thresholds and both tie methods, the proportion of subnetworks that were intransitive increased with the size of the subnetwork. For instance, with a 50% threshold and the conservative DAG approach, 0%, 0.1%, and 0.2% of the two-, three-, and four-species subnetworks, respectively, were not DAGs, while with the same threshold and the liberal DAG approach, these percentages were 3.6%, 4.5%, and 5.6%, respectively. When we increased the threshold with the liberal DAG approach, a larger proportion of subnetworks were no longer DAGs, while the opposite was true of the conservative DAG approach (Fig. 4). This is expected, given that tied interactions are converted to bidirectional edges with the liberal approach, which by definition imposes cycles in the subnetwork.

**Figure 4.**
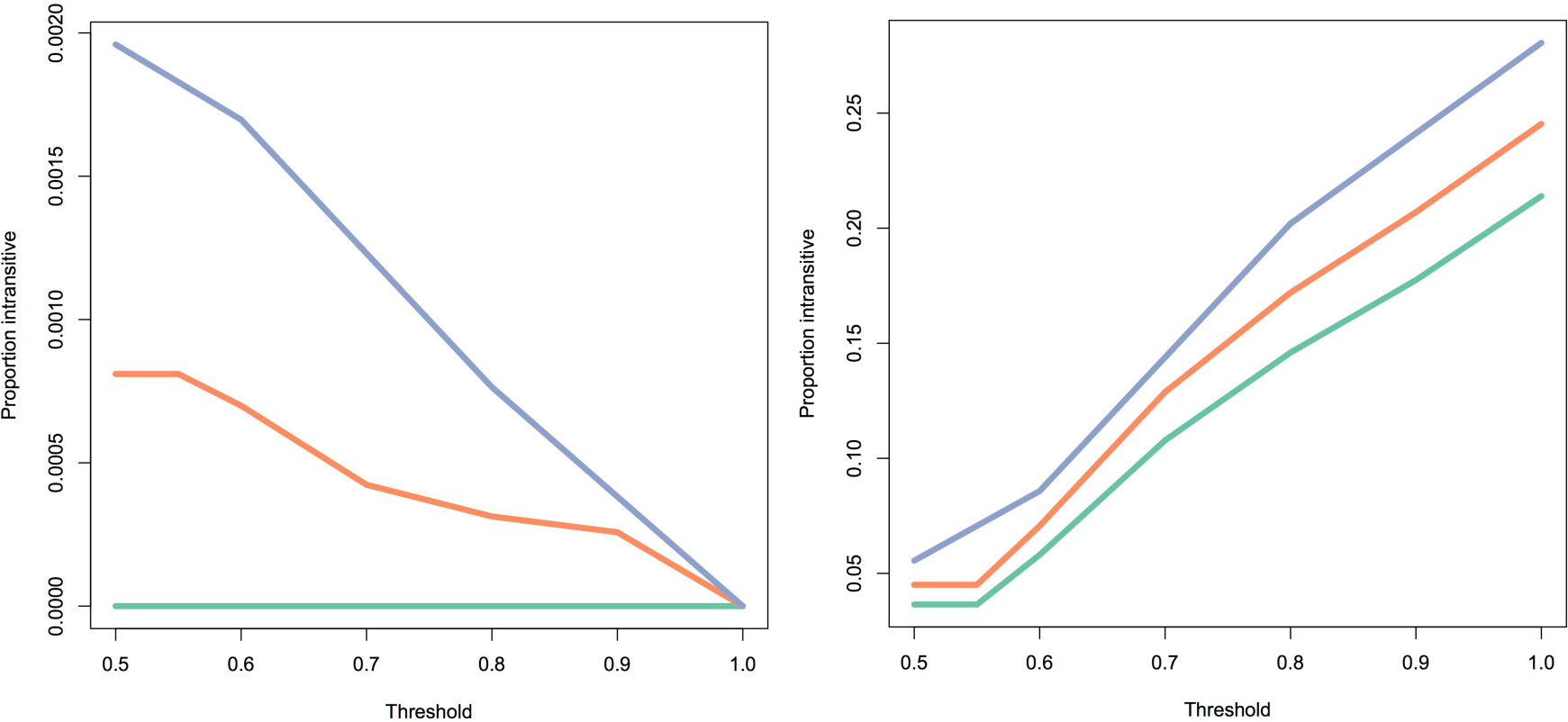
Illustration of how proportion of subnetworks that are intransitive varies as a function of threshold and method of handling ties. Green, orange and blue lines correspond to two-three- and four-species subnetworks, respectively. With the conservative DAG approach (left panel), pairwise interactions where neither species won more than the threshold are removed, and intransitivity therefore decreases as the threshold increases. With the liberal DAG approach (right panel), pairwise interactions below the threshold are set to bidirectional arrows. The proportion of subnetworks that are intransitive therefore increases when the threshold is increased with this method, with the caveat (Fig. 1) that not all networks identified as cyclic are truly intransitive here.

These tests of intransitivity show that while the network approximated a linear dominance hierarchy, it was not perfectly so; even at the pairwise level, 21% of interactions exhibited variation in directionality (as assessed with the 100% threshold, liberal DAG method). Results from our null model simulations with the 50% threshold, conservative DAG approach showed that House Finch, Red-bellied Woodpecker (*Melanerpes carolinus*), and Blue Jay (*Cyanocitta cristata*) were embroiled in the fewest cyclic relationships, but that overall, almost all species had negative SES values, indicating that most triads were transitive. Only the poorly sampled Red-headed Woodpecker (*M. erythrocephalus*), Townsend’s Warbler (*Setophaga townsendi*) and Hermit Thrush (*Catharus guttatus*) had positive SES values. Finally, species infrequently observed at feeders (kinglets, thrushes, warblers, creepers, etc.) tended to most frequently be involved in intransitive relationships (Table S1). In summary, the dominance hierarchy approximated a linear pecking order as assessed at the triad level, although our ability to explore the extent of intransitivity, specifically intransitivity within large subnetworks, was hampered by computational limitations.

## DISCUSSION

In this paper we used a large database of citizen-scientist-observed behavioral interactions to derive an interspecific social dominance hierarchy of North American feeder birds (Fig. 2). This hierarchy concurred with published information at the species-pair level (Rodewald 2015). Despite observations encompassing an ecologically wide range of birds across the breadth of the North American continent, the resulting hierarchy was nearly linear. Whether assessed with the triangle transitivity method (Shizuka and McDonald 2012) or the directed acyclic graph (DAG) method introduced here, most interactions were transitive. Thus, the continental dominance hierarchy of feeder birds, as determined by citizen scientists, appeared to deviate only slightly from a linear pecking order; near linear hierarchies are frequently observed in biology (Landau 1951; Chase 1982). This result sets up a number of interesting questions. For example, can the presence of a similarly ranked species preclude another’s occurrence, and is there geographic or temporal variation in a species’ ranking?

Species’ positions in the dominance hierarchy were related to body mass, although certain lineages were more or less dominant than expected for their mass (e.g., chickadees and titmice, Fig. 3). Martin and Ghalambor (2014) suggested that morphological adaptations for foraging on trunks might reduce performance in aggressive encounters. This suggestion was based on the inclusion of woodcreepers (Furnariidae) in their study, which did not include woodpeckers. Our results, which do not include woodcreepers, show woodpeckers to be more dominant than expected based on their body mass. Thus, trunk foraging *per se* does not appear to explain these clade-level differences. Alternatively, a woodpecker can sustain impressive forces on its skull without suffering brain injury (Wang et al. 2011), a morphological character that seems relevant for an animal that fights in large part with its head. Identifying additional morphological or behavioral characteristics that explain variance in dominance, beyond body mass, awaits future study. In addition to these large shifts in dominance towards the root of the phylogeny, leading certain clades to deviate from the overall mass-to-dominance relationship, there were also a number of large shifts in dominance towards the tips of the phylogeny (hence, the low lambda from the PGLS regression). For instance, Canyon Towhee (*Melozone fusca*) was inferred to be notably more dominant than the closely related, similarly sized California Towhee (*M. crissalis*). Both species were infrequently observed by participants (Table S1), so this apparently large shift in dominance may simply be a function of limited information. Confirming these patterns, and identifying which factors explain these large shifts in dominance, both between higher taxonomic levels and between close relatives, requires further data collection and analyses.

A large body of theoretical literature supports the notion that intransitivity in dominance relationships promotes species coexistence (Laird et al. 2006; Laird and Schamp 2008; Allesina and Levine 2011). This has been empirically demonstrated in the lab (Kerr et al. 2002), and it has received some attention in plant competition literature (Keddy and Shipley 1989; Soliveres et al. 2015), but empirical tests of this idea are infrequent, particularly in animal systems. Overall, we found few well-supported, ecologically compelling intransitive relationships at the three- and four-species scale. Of those intransitivities of possible ecological relevance, we found the House Finch, Purple Finch (*Haemorhous purpureus*), and Dark-eyed Junco (*Junco hyemalis*) subnetwork to be intransitive, with the House Finch dominant to the Purple Finch (*n*=8), the Purple Finch dominant to the junco (*n*=3), and the junco dominant to the House Finch (*n*=14, and 7 observations of the opposite interaction). If this holds given further observations, we would predict increased negative consequences of Dark-eyed Junco-House Finch competition in areas such as the southwestern USA, where Purple Finches do not regularly occur.

Most of the intransitive relationships we detected were only supported by small sample sizes. Indeed, when we removed all single interactions (i.e., those where species A had only interacted once with B), and summarized the network with the 75% conservative DAG method, the entire dominance hierarchy was transitive. Moreover, multiple intransitive relationships might be driven by a single odd observation, i.e. knock-on effects. For instance, a number of intransitive relationships were rooted in a single observation of a White-throated Sparrow (*Zonotrichia albicollis*) interacting with a European Starling (*Sturnus vulgaris*), an interaction where the former displaced the latter. It seems likely this relationship, and perhaps others, will be reversed given more data, and the degree to which these intransitive relationships will appear more linear with added data remains to be seen. We did find well-supported instances of shifting dominance relationships, but these were not truly intransitive relationships. For instance, both White-breasted Nuthatch and Tufted Titmice (*Baeolophus bicolor*) tended to displace Black-capped Chickadee (*Poecile atricapillus*), but the nuthatch and titmouse displaced one another at approximately equal rates, a fact that has been noted before (Waite and Grubb 1988), and may be related to sexually associated intraspecific differences in dominance (i.e., male titmice may be dominant to female nuthatches).

Numerical measures of intransitivity are complicated by the rules used to identify and deal with ties. In general, transitivity decreases when ties are converted to bidirectional edges, while it increases when these ties are removed from the network; transitivity also decreases as larger numbers of species are considered in subnetwork of the broader dominance hierarchy. Our DAG method differs slightly from the triangle transitivity in that the former considers all possible subnetworks of a given size, while the latter only considers triads where all three relationships are known. Regardless of the method used, the result is qualitatively the same, the hierarchy was mostly linear as assessed at the computationally accessible scale of three- and four-species subnetworks. Certainly, diversity promoting intransitivities may be present in the dominance hierarchy of North American, but we are currently limited in our ability to identify them.

As we know of no exercise comparable in geographic scope, we use this opportunity to ask whether the definition of a continent-wide dominance hierarchy is biologically meaningful. For example, given that a Black-billed Magpie (*Pica hudsonia*) in Alberta and a Brown-headed Nuthatch (*Sitta pusilla*) in South Carolina stand very little chance of directly interacting, is it reasonable to rank them on the same scale? As stated above, on a pairwise level, our linear dominance hierarchy was in close accord with published examples (Rodewald 2015). It is notable that this consistency emerged from our hierarchy even when we did not have observations of direct interactions between the close relatives in question. It is possible to rank species in a corporative framework, where some species hold equivalent ranks (Fushing et al. 2011), but such approaches have not yet been implemented in available software. Thus, while there is room for improvement in the algorithm used to assign rank, and as more data are contributed species’ precise positions in the hierarchy may shift slightly, our continental dominance hierarchy appears to be biologically reasonable. Moreover, its structure is interesting in its own right, and it can be brought to bear on a wide variety of research questions. For example, as no-analog communities (Jackson and Overpeck 2000) are generated as a product of climate change, species introductions, and other anthropogenic activities, interspecific dominance hierarchies linking currently geographically separated taxa may actually provide a predictive framework for understanding which species will and will not be able to coexist.

Social dominance interactions like those studied here have well-documented consequences on species’ acquisition of resources (e.g., Samuels et al. 1984). Even within our study, dominant species clearly had preferential access to food at feeders. However, whether a link exists between social dominance interactions and competitive dominance over longer evolutionary timescales is unknown. Certainly, these interactions shape species’ behaviors indirectly. For instance, species may shift their foraging times and locations to avoid close competitors that they are unable to exclude. As an example, less dominant hummingbirds may feed on lower-quality nectar patches (Wolf et al. 1976). Behavioral interactions at a local scale may therefore scale up to larger phenomena on continental scales, including migration timing and distance (Freshwater et al. 2014), and such interactions may even play a role in shaping species’ distributions on a continental scale (Robinson and Terborgh 1995; Pasch et al. 2013). These biological hypotheses can be examined only by describing dominance relationships across communities of birds over very large regions. By partnering with an existing citizen science project, we provided the first continental-scale view of these local, infrequently observed interactions at communal food resources. We see our work, where a targeted behavioral study was linked with an existing citizen science project, as a model for gathering robust natural history data at broad spatiotemporal scales.

## ACKNOWLEDGEMENTS

We thank the citizen scientists of Project FeederWatch who participated in the interactions project; Anne Marie Johnson, Chelsea Benson, Kerrie Wilcox, and Lisa Larson for facilitating the link between this project and Project FeederWatch; the Harmon lab for statistical input; and G. Leighton, N. Mason, D. Toews, J. Berv, V. Rohwer, J. Walsh and R. Laird for helpful discussion and comments on earlier versions of the manuscript.

## SUPPLEMENTARY TABLE LEGEND

Table S1. Summary of sample sizes used to create the continental dominance hierarchy. The first column details the absolute number of observations per species included in the study, while the second summarizes the number of unique households that submitted interactions that included that species. The third column provides four-letter AOU codes. These codes are used to identify species in Fig. 2.

